# Construction and characterization of coronavirus nonstructural protein 3-host protein interaction networks unravel an important role of cleavage and polyadenylation specificity factor 6 in regulation of viral RNA replication

**DOI:** 10.1101/2025.02.27.640555

**Authors:** Xinxin Sun, Lixia Yuan, Zihong Hu, Yuzhu Lai, Bei Yang, Jiawen He, Rui Ai Chen, Ding Xiang Liu

**Affiliations:** Integrative Microbiology Research Centre, South China Agricultural University, Guangzhou 510642, China; Zhaoqing Branch Center of Guangdong Laboratory of Lingnan Modern Agricultural Science and Technology, Zhaoqing 526238, China; College of Veterinary Medicine, South China Agricultural University, Guangzhou 510642, China; Key Laboratory of Viral Pathogenesis & Infection Prevention and Control (Jinan University), Ministry of Education, Guangzhou, 510632, China

**Keywords:** Coronavirus, SARS-CoV-2, IBV, Nsp3, CPSF6, Protein interactomes, gRNA and sgRNA, Viral replication

## Abstract

Coronavirus nonstructural protein 3 (nsp3) plays a crucial role in viral replication and immune evasion. However, functional and proteomic characterization of this protein, especially the interaction networks between nsp3 from different coronaviruses and host cell factors, is hindered by its huge size, complex structural feature and the presence of multiple transmembrane domains. In this study, we report the application of a high-performance cytoplasmic expression system to efficiently and accurately express the full-length nsp3 from betacoronavirus severe acute respiratory syndrome coronavirus 2 (SARS-CoV-2) and gammacoronavirus infectious bronchitis virus (IBV), to investigate their interactions with host proteins using proteomics approaches. Our study identified 1,150 host proteins that interact with IBV nsp3 and 920 with SARS-CoV-2 nsp3. Among them, 658 are shared by the two nsp3 proteins. Further validation and preliminary characterization of seven selected candidates, DDX5, DDX39, DHX9, elF4A3, SRRT and CPSF6, demonstrated the reproducibility and reliability of the proteomics data. More interestingly, an important regulatory role of the nsp3-CPSF6 interaction in the replication and transcription of IBV gRNA and sgRNA was unraveled. The construction of nsp3-host protein interaction networks from two distantly related coronaviruses would have provided a foundation for future studies of host cell factors in the regulation of coronavirus replication and pathogenesis.

## 1. Introduction

Coronaviruses (CoVs) cause highly infectious and prevalent respiratory and enteric diseases in humans and animals, imposing immense public health concerns and economic losses worldwide. CoVs are divided into four genera, *Alphacoronavirus*, *Betacoronavirus*, *Gammacoronavirus*, and *Deltacoronavirus*, belonging to a large subfamily, *Orthocoronavirinae*, of the *Coronaviridae* family within the *Nidovirales* order [1]. Common human and animal pathogens include the novel human coronavirus severe acute respiratory syndrome coronavirus 2 (SARS-CoV-2), a *betacoronavirus*, avian infectious bronchitis virus (IBV), a *gammacoronavirus* infecting chicken, and porcine epidemic diarrhea virus (PEDV), a member of the *alphacoronavirus* genus.

CoV genome is a single-stranded, positive sense RNA, with a size of 27∼32 kilobases [2]. The genomic RNA (gRNA) contains a 5’ cap structure, a 3’ poly A tail, and multiple open reading frames (ORFs), with the capacity to encode 4 structural proteins, spike (S) protein, envelope (E) protein, membrane (M) protein, nucleocapsid (N) protein, 15-16 nonstructural proteins (nsps), and several accessory proteins [2]. These proteins are translated from the gRNA and a set of nested positive-strand subgenomic RNAs (sgRNAs), due to the presence of a number of transcriptional regulatory sequences (TRS) in the gRNA. Viral RNA replication begins with transcription of the full-length negative-strand gRNA (-gRNA) and a set of nested negative-strand sgRNAs (-sgRNAs) from the 3’ end of gRNA by the replication-transcription complex (RTC) using continuous and discontinuous transcription mechanisms, respectively [2]. These -gRNA and -sgRNAs then serve as templates for the production of new gRNAs and sgRNA [2,3].

Among the 15-16 nsps, the largest is the multidomain nsp3, with an average molecular weight of approximately 200 kDa and containing various enzymatic activities essential for viral replication [4]. As a crucial component of RTC, nsp3 induces the rearrangement of cellular membranes to form double-membrane vesicles (DMVs) or spherules along with nsp4 and nsp6, and assembles into RTC [5–7]. In addition to its role as a PLpro that cleaves nsp1–nsp3 from the polyproteins, nsp3 also has deubiquitination (DUB) and de-ISGylation activities, serving interferon (IFN) antagonism functions [8]. The macrodomain of nsp3 also involves ADP-ribose in regulating the innate immune response [9]. The huge size and multifunctional nature of nsp3 poses an interesting but challenging question on how to efficiently and accurately determine host proteins that may interact with it and modulate its functions. During the global COVID-19 pandemic, 332 high confidence SARS-CoV-2 host protein interactions (PPIs) were first identified using AP-MS after overexpression of 26 viral proteins, excluding nsp3 and nsp16 [10]. Subsequently, Xu et al. mapped the interaction network among RTC-related viral proteins using a novel protein interaction technology called Compartmentalization of Protein-Protein Interactions in Cells (CoPIC) [11]. However, due to the challenge in efficient expression of the full-length nsp3, the nsp3-host protein interactome is less reported. Most studies used the strategy to express nsp3 segments to determine the interacting proteins utilizing AP-MS [12]. A more recent study successfully overexpressed the full-length SARS-CoV and SARS-CoV-2 nsp3 proteins and used multi-omics methods to investigate the unique and shared functions of these nsp3 proteins [13].

We have previously reported the development of a high-performance expression system to express the full-length nsp3 from IBV and SARS-CoV-2 [14]. In this study, we report the nsp3-host protein interactomes in cells overexpressing IBV and SARS-CoV-2 nsp3 using this expression system, validation of seven selected common nsp3-interactors, including DEAD-box RNA helicase 5 (DDX5), DExD-box helicase 39B (DDX39), DExH-Box helicase 9 (DHX9), eukaryotic initiation factor 4A3 (elF4A3), Serrate RNA effector molecule homolog (SRRT) and cleavage and polyadenylation specificity factor subunit 6 (CPSF6). Further preliminary functional characterization of the interaction between IBV nsp3 and CPSF6 in regulation of the replication of IBV and human coronavirus OC43 (HCoV-OC43). This study presents a comprehensive interaction network of nsp3 from two coronaviruses in different genera and would have provided a solid basis for further elucidation of the involvement of host proteins in coronavirus replication and pathogenesis.

## 2. Materials and methods

### 2.1 Viruses, Cells and Antibodies

Vero adaptive IBV (p65) was obtained from the American Type Culture Collection (ATCC) and propagated in Vero cells for 65 passages, as previously described [15,16]. rIBV-HiBiT was rescued using reverse genetics technology by insertion of an 11-amino acid HiBiT tag into the S protein [17]. All viruses were propagated and passaged in Vero cells. H1299 and HEK293T cells were cultured in Dulbecco’s Modified Eagle Medium (DMEM, Gibco) containing 10% fetal bovine serum (FBS) and 1% penicillin-streptomycin (Gibco) at 37°C and 5% CO₂. In the infection experiments, cells were treated as previously described [14], and then washed with PBS before being infected with IBV at a multiplicity of infectivity (MOI) of approximately 2 and incubated in serum-free DMEM. After 2 h of absorption, cells were washed twice with PBS to remove the unbound viruses.

Antibodies against FLAG, HA and β-actin were bought from TansGen biotech (Beijing, China), against CPSF6 were bought from Proteintech (Wuhan, China), and antibodies against HCoV-OC43 N and HiBiT tag were purchased from SinoBiological (Beijing, China) and Promega (Wisconsin, USA), respectively. Anti-IBV N protein serum was made by immunization of rabbits as previously described [14].

### 2.2 Immunoprecipitation and Mass Spectrometry

Immunoprecipitation was carried out essential as previously described [14]. Briefly, cells transfected with appropriate plasmid DNA or infected with viruses were lysed with RIPA lysis buffer containing protease inhibitors by thorough mixing and then on a table rotator at 4°C for 15 min. After clarifying by centrifugation at 10,600× g for 10 min, the supernatant was transferred to a new centrifuge tube containing pre-washed beads, followed by adding lysis buffer to a final volume of 1 mL plus 10 μL of protease inhibitors, and incubation at 4°C for 2 h. The bound beads were then collected by centrifugation at 8,300× g, 4°C for 40 sec, washed with 1 mL of lysis buffer three times, and analyzed by Western blot.

Nsp3 enrichment was performed on HEK293T cells 24 h post-transfection using affinity gels with the corresponding antibody tag. The resulting protein samples were sent to Sangon Biotech for mass spectrometry identification, and the results were analyzed using the Mascot protein database. After excluding nonspecific proteins from the control group, the UniProt database was used to search for information on host proteins interacting with IBV and SARS-CoV-2 nsp3 proteins, providing data for the next steps in proteomic analysis.

### 2.3 RNA extraction and RT-PCR Analysis

Cells were lysed with 1 mL TRIzol and 200 μL of RNA extraction agent provided, and the aqueous phase was collected by centrifugation at 12,000 × g at 4°C for 15 min. Total RNA in the aqueous phase was precipitated by mixing with an equal volume of isopropanol and centrifugation at 12,000 × g at 4°C for 15 min, washed twice with 70% ethanol, and dissolved in 30 μL RNase-free water. RT-PCR was performed by mixing 2 μg of total RNA with 1 μL Oligo(dT) (10nM) and 1 μL dNTP Mixture (10mM) in a 10 μL reaction mixture, incubation at 65°C for 5 min and on ice for 2 min, followed by adding 5X Reverse Transcriptase Buffer, RNase Inhibitors, and Reverse Transcriptase M-MLV, and incubating at 42°C for 60 min. The enzyme was then inactivated by incubation at 72°C for 15 min.

The relative expression levels of IBV and OC43 gRNA and sgRNA were measured by quantitative real-time PCR (RT-qPCR) using the Talent qPCR PreMix SYBR Green kit (Tiangen). The nucleotide sequences of forward and reverse primers used in this study were listed as follows. IBV gRNA 5’GTTCTCGCATAAGGTCGGCTA3’ and 5’GCTCACTAAACACCACCAGAAC3’, IBV sgRNA2 5’GCCTTGCGCTAGATTTTTAACTG3’ and 5’AGTGCACACAAAAGAGTCA CTA3’, IBV sgRNA4 5’GCCTTGCGCTAGATTTTTAACTT3’ and 5’GTACAATTTGTCTCGTT GGGC3’, IBV sgRNA6 5’GCCTTGCGCTAGATTTTTAACTT3’ and 5’GCTTTACCGCTTGCC ATG3’, HCoV-OC43 gRNA 5’TGCGAACAAGACTCGTGTGAT3’ and 5’ACACCTGAAGATTGT GCCAT3’, HCoV-OC43 sgRNA2 5’CGGATGCAGCACTGTCCATT3’ and 5’AAGTCTAGCCG GTGATGAGG3’, HCoV-OC43 sgRNA4 5’AATTTGTGCACTCCTGATCCT3’ and 5’AACAGC AAGACCCGAACAGT3’, and HCoV-OC43 sgRNA6 5’AGTGGCATTTTTGGCAACTTTT3’ and 5’TCCACATCAAGGACTGGTGG3’.

The nucleotide sequences and primer pairs used for PCR amplification of host cell genes in H1299 and Vero cells are GAPDH 5’CTGGGCTACACTGAGCACC3’ and 5’AAGTGGTCGTTGAGGGCAATG3’, DDX5 5’ACAGTTAGAGGTCACAACTGC3’ and 5’TCCCTGAGCTTGAATAGCAGT3’, DDX39B 5’TCCAGGCCGTATCCTAGCC3’ and 5’GCATGTCGAGCTGTTCAAGC3’, DHX9 5’CAGGAGAGAGAGTTACTGCCT3’ and 5’CTCTGCTGCTCGGTCATTCTG3’, CPSF6 5’CCTGGTGGGGACAGATTTC3’ and 5’GTGGTGGAGTCTGTCCAGC3’, eIF4A3 5’GGGGCATCTACGCTTACGG3’ and 5’TCTCTTGTGGGAGCCAAGATC3’, SRRT 5’CACCATGTCCTGCCTATCCAG3’ and 5’CACTCCTCATCTTTGTGCGC3’, and UBA1 5’TCGCCGCTGTCCAAGAAAC3’ and 5’AGTAAAGGCCCTCGTCTATGTC3’.

### 2.4 Plasmid Construction, Transfection and Western Blot Analysis

Plasmids K1E-HibiT-SARS-CoV-2-nsp3 and K1E-HibiT-IBV-nsp3 were cloned as previously described [14]. DDX5, DDX39B, DHX9, elF4A, SRRT, UBA1 and CPSF6 were amplified from 293T cells by RT-PCR, cloned into pXJ40-Flag by homologous recombination with corresponding primer pairs, and verified by nucleotide sequencing.

Transfection of either plasmid DNA or siRNA into HEK293T cells was carried out as previously described [14]. Briefly, plasmid DNA and TransIntro EL reagent were diluted in Opti-MEM (Gibco), and the transfection mixture was added to cells after incubation at room temperature for 20 min. Cells were further incubated in Opti-MEM supplemented with 10% FBS and collected at indicated time points, total cell lysates were prepared and clarified by centrifugation. Proteins were separated by sodium dodecyl sulfate–polyacrylamide gel electrophoresis (SDS-PAGE), transferred to a nitrocellulose membrane (BIORAD), and detected by Wesern blot with appropriate primary and secondary antibodies.

The HiBiT-tagged proteins were detected using the Nano-Glo HiBiT blotting system (Promega) as previously described [17], and the chemoluminescence of the HiBiT-tagged protein was detected using the Amersham ImageQuant™ 800 (Cytiva).

### 2.5 Immunofluorescence staining

Cells were subjected to immunofluorescence staining as previously described [18]. Briefly, H1299 cells seeded on 96-well plates were infected rIBV-HA-3541 (at a MOI of 2) for 24 h fixed with 4% paraformaldehyde for 20 min, permeabilized by treatment with 0.25% Triton X-100 in phosphate-buffered saline (PBS) for 10 min, washed three times with PBS, and blocked with PBS containing 1% bovine serum albumin (BSA) plus Tween 20 (0.1% [vol/vol]) for 2 h at room temperature. Cells were then incubated with the primary anti-CPSF6 and HA antibodies, respectively, at 4℃ overnight and with the secondary antibody for 2 h at room temperature, washed with PBS three times, nuclear stained with 49,6-diamidino-2-phenylindole (DAPI). Images were obtained with a fluorescence microscope.

### 2.6 Statistical analysis

A one-way analysis of variance (ANOVA) was conducted to assess significant differences between the specified samples and their corresponding control samples. The significance levels are indicated by the P value (ns, nonsignificant; *, P, 0.05; **, P, 0.01; ***, P, 0.001).

## 3. Result

### 3.1 Proteomic analysis of nsp3-host protein interactomes

To investigate the host proteins that interact with nsp3 in coronavirus-infected cells and their functional roles, we conducted an affinity mass spectrometry on nsp3-expressing cells to analyze the potential interactions between host proteins and nsp3 from IBV and SARS-CoV-2. Plasmids pK1E-HA-SARS-CoV-2-nsp3, pK1E-FLAG-IBV-nsp3, carrying HA tagged SARS-CoV-2 nsp3 and FLAG-tagged IBV nsp3, respectively, and their corresponding empty vectors were transfected into HEK293T cells (Figure 1A). The potential nsp3-interacting proteins were enriched using affinity gel conjugated with the appropriately labeled antibody at 24 h post-transfection (Figure 1A). The collected samples were split into two aliquots and incubated with same amounts of affinity gel beads. The first aliquot was subjected to Coomassie dye staining following SDS-PAGE to validate IP efficiency and to determine the presence of multiple host proteins that may interact with nsp3 (Figure 1B). The second aliquot was sent to a commercial laboratory for mass spectrometry analysis (Figure 1A), and data obtained were analyzed using Mascot software. After removing the peptide segments identified from the non-specific elution of the empty vector control group, the data were searched in the UniProt database to gather relevant protein information. As shown in Figure 1C, 1150 host proteins interacted with IBV nsp3, and 920 with SARS-CoV-2 nsp3, with a difference of 170, were identified (Supplementary Table S1A, S1B). Among them, 658 were shown to interact with both IBV nsp3 and SARS-CoV-2 nsp3 (Figure 1C). These results indicate that coronavirus nsp3 may interact with a multitude of common host proteins to facilitate their replication during their infection cycles, but different coronavirus nsp3 may also recruit sets of specific host proteins to play unique roles in viral pathogenicity and tissue/cell tropism.

**Figure 1.**
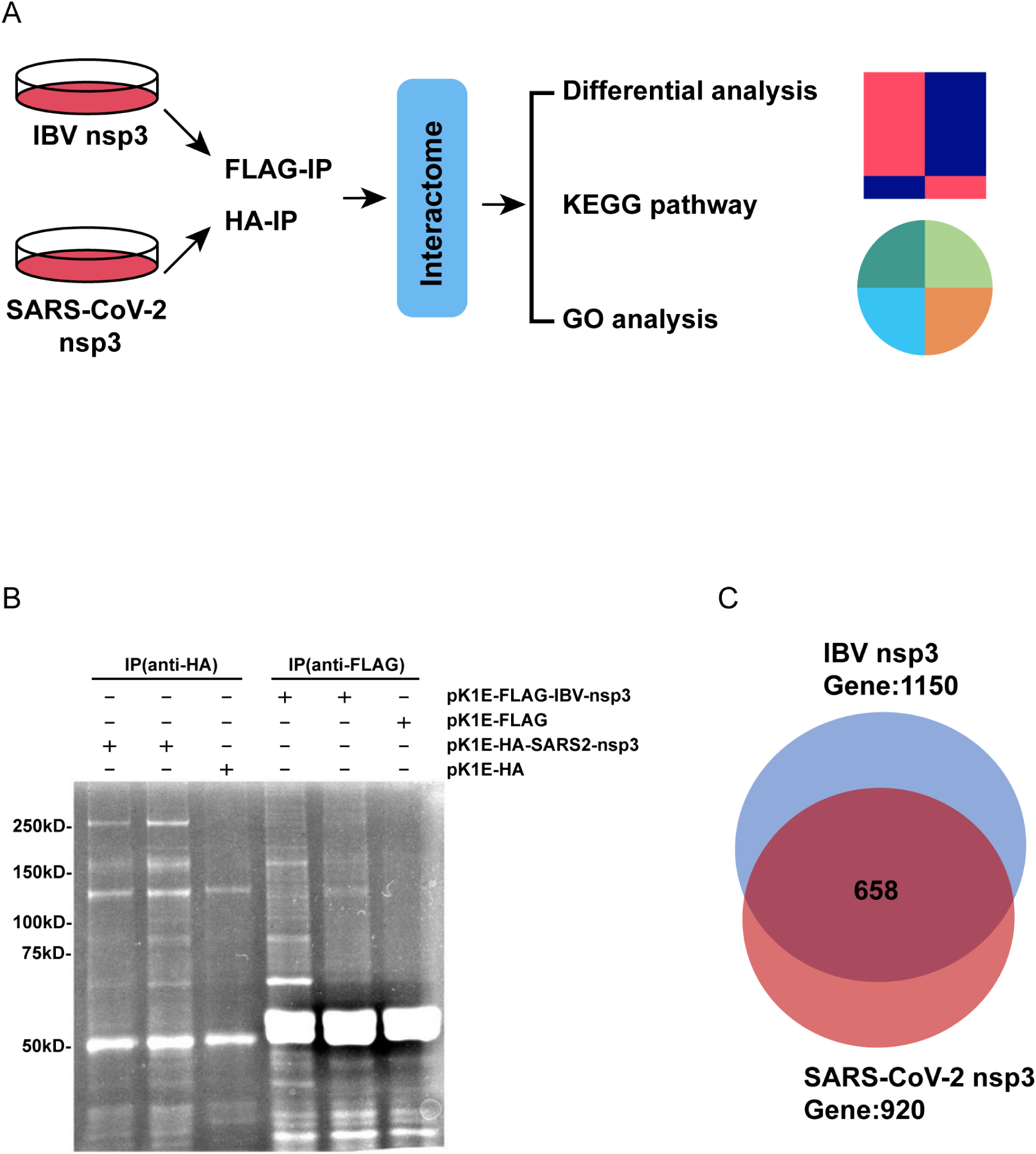
Identification of interacting proteins using AP-MS. (A) AP-MS workflow to identify host proteins interacted with nsp3 from IBV and SARS-CoV-2. HEK293T cells were co-transfected with pK1E-FLAG/HA and either pK1E-FLAG-IBV or pK1E-HA-SARS2-nsp3, lysates were prepared and immuno-purified using anti-FLAG or HA beads to enrich nsp3 in complex with host interactors. The interacting proteins were identified by MS and further analyzed by GO and KEGG analyses. (B) SDS-PAGE analysis of the overexpression IBV and SARS-CoV-2 nsp3. HEK293T cells were co-transfected as described in (A), and harvested at the indicated time points. Lysates were prepared and incubated with the respective affinity gel with the related labeled antibody. The proteins were separated by SDS-PAGE and visualized by staining with Coomassie dye. Cells transfected with pk1E-FLAG/HA were used as a negative control. (C) Numbers of interacting proteins in IBV and SARS-CoV-2 nsp3. Venn diagram shows the numbers of unique and shared potential host proteins interacting with nsp3 from IBV and SARS-CoV-2, as indicated.

### 3.2 Gene Ontology (GO) enrichment analysis and Kyoto Encyclopedia of Genes and Genomes (KEGG) Pathway enrichment analysis of nsp3-interacting proteins

GO analysis of the identified interacting proteins summarized 10 items with significant differences in the enrichment analysis of molecular biological processes, as illustrated in Figure 2. Among IBV nsp3-interacting proteins, the top five biological processes identified through this analysis are RNA metabolism, protein translation, biogenesis of nucleoprotein complexes, regulation of the cell cycle, and regulation of mRNA metabolism (Figure 2A). For SARS-CoV-2 nsp3-interacting proteins, the primary processes include protein translation, RNA metabolism, biogenesis of ribonucleoprotein complexes, neutrophil regulation, and RNA splicing regulation (Figure 2B). Although nsp3 from these two coronaviruses would share similar functions in viral replication, they may also play unique roles in viral pathogenicity and virus-host interaction. By interacting with host proteins predominantly in these biological processing, this viral non-structural protein would render significant influence on RNA metabolism and protein translation within host cells, ultimately facilitating viral replication as well as regulating cellular processes.

**Figure 2.**
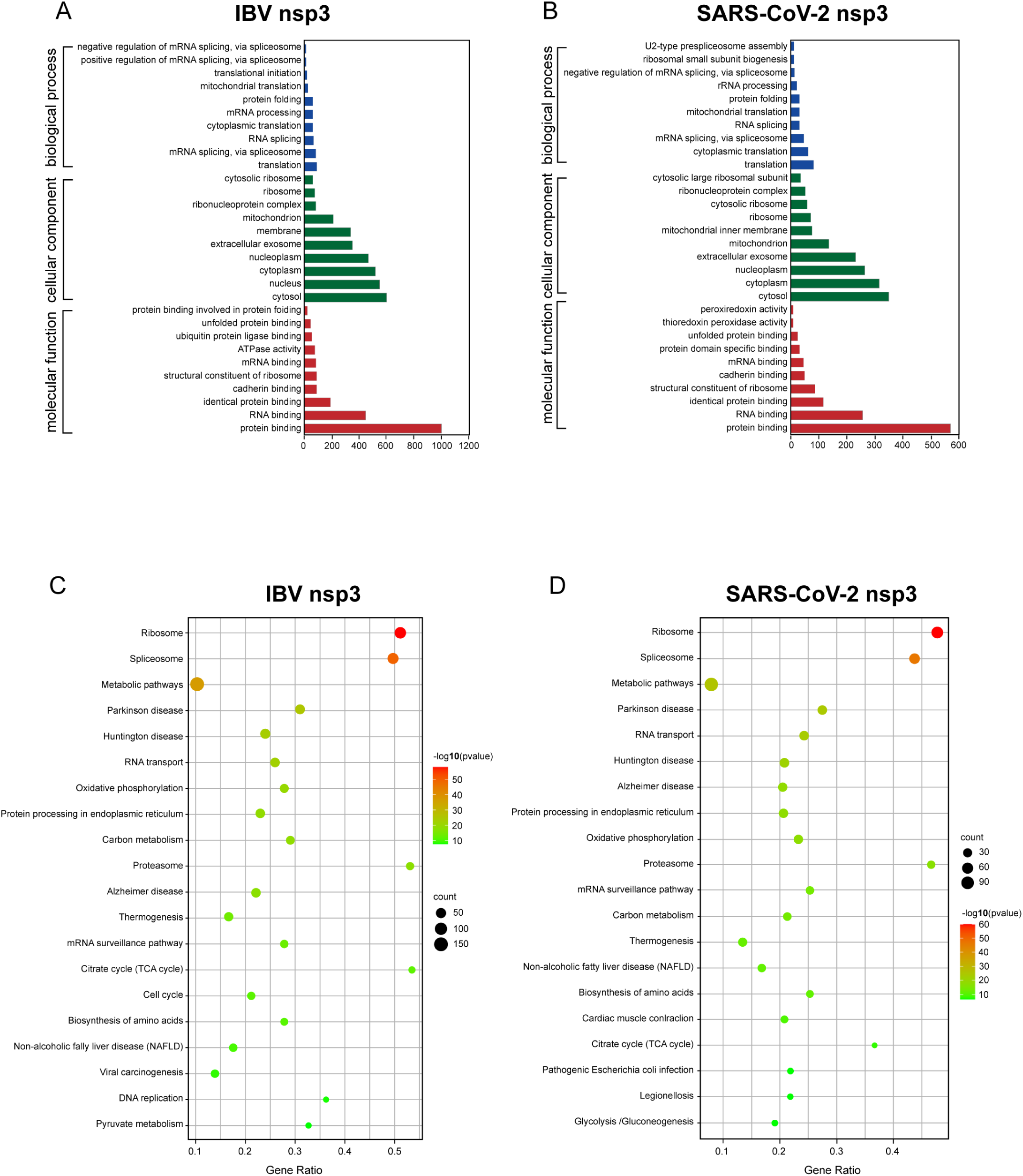
GO enrichment and KEGG pathway analysis of nsp3-interacting proteins. (A/B) GO enrichment analysis of host protein interacting with nsp3 from IBV (A) and SARS-CoV-2 (B). The vertical coordinate represents the GO enrichment item, and the horizontal coordinate represents the number of interacting protein genes under this item. (C/D) The KEGG pathway enrichment analysis of host proteins interacting with nsp3 from IBV (C) and SARS-CoV-2 (D). The vertical coordinate is the KEGG pathway, and the abscissa is the ratio of interacting protein genes to all genes in the pathway. The dot size represents the number of genes, and the darkness of the dot color represents the significance Q value (the darker the color, the smaller the significance Q value, and more reliable the data).

KEGG analysis of the genes that interact with these two nsp3 was then conducted, showing that proteins interacting with IBV nsp3 were involved in 294 pathways, while those interacting with SARS-CoV-2 nsp3 were involved in 269 pathways. Among the top 20 pathways with the smallest significant Q values for summary analysis, 16 were identically shared with IBV and SARS-CoV-2 nsp3. These common pathways include ribosome, spliceosome, metabolic pathways, Parkinson’s disease, Huntington’s disease, RNA transport, oxidative phosphorylation, protein processing in the endoplasmic reticulum (ER), carboxylation and carbon metabolism, proteasome, Alzheimer’s disease, thermogenesis, mRNA surveillance pathway, citrate cycle, biosynthesis of amino acids, and non-alcoholic fatty liver disease (Figure 2C and 2D). Once again, most of these enrichment pathways are related to mRNA metabolism and protein synthesis, in addition to some pathways participating in disease and metabolism. This suggests that coronavirus nsp3 may influence cell metabolism and subsequently regulate viral replication mainly by binding to host proteins involved in the mRNA maturation, splicing, transport, and protein translation and synthesis.

### 3.3 Statistical analysis and mapping of nsp3-host protein interactomes

All the interacting proteins were further reordered, and the top 50 proteins with the highest MS intensity in the IBV nsp3-interacting proteins were illustrated in Figure 3A. Among them, Nucleophosmin 1 (NPM1), a nucleolar chaperone involved in diverse biological functions, including genomic stability and tumorigenesis, ribosome biogenesis, centrosome duplication, DNA repair, and response to cellular stress, displays the highest MS intensity [19]. Statistically the KEGG pathway analysis revealed that seven out of the top 50 proteins were enriched in the spliceosome pathway (Figure 3B).

**Figure 3.**
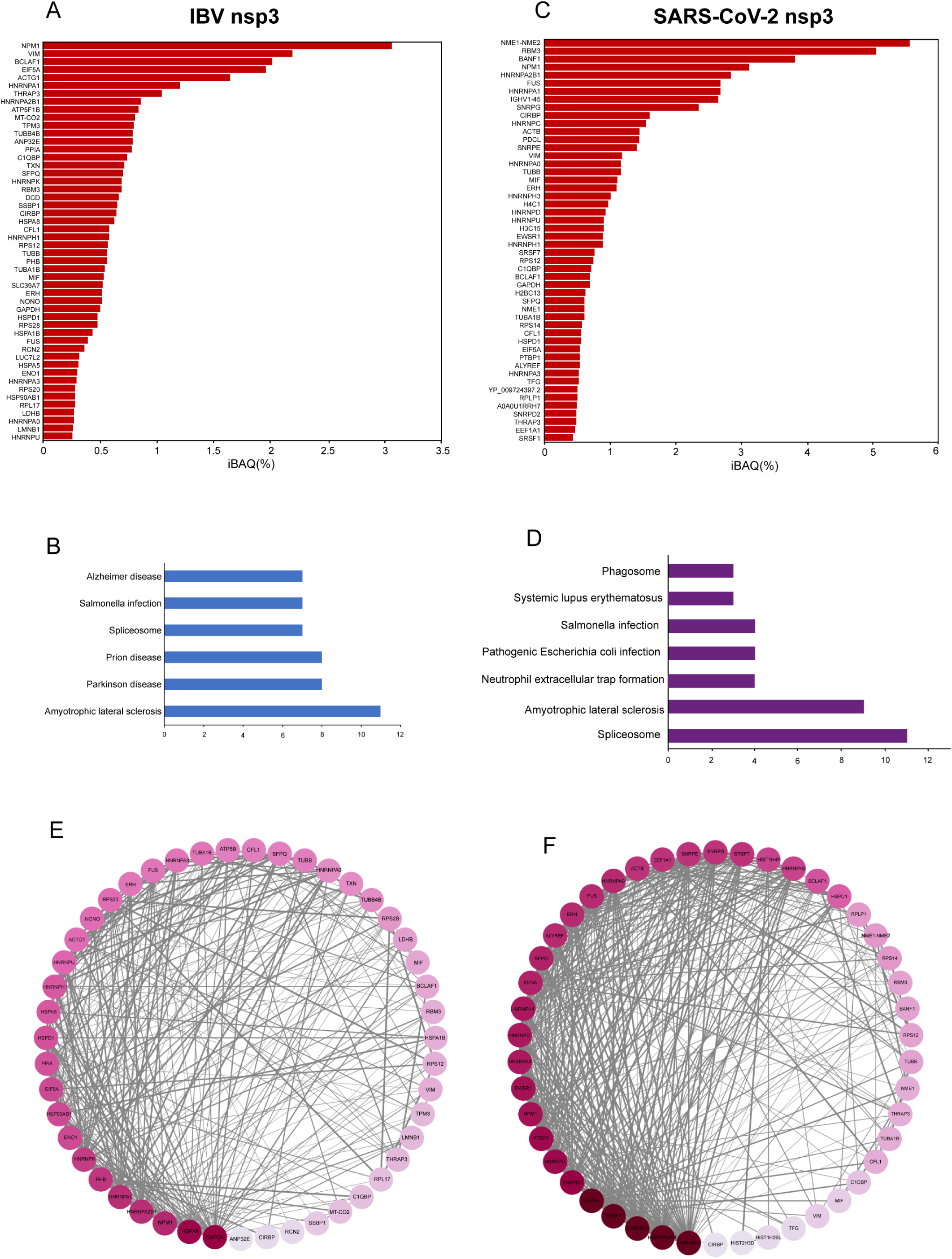
Statistical analysis of the top 50 proteins interacting with nsp3. (A) &(D) MS intensity of host protein interacting with nsp3 from IBV (A) and SARS-CoV-2 (D). The vertical axis represents the names of the genes corresponding to the proteins, and the horizontal axis displays the corrected strength value (iBAQ%). (B) &(E) KEGG analysis of top 50 host proteins interacting with nsp3 from IBV (B) and SARS-CoV-2 (E), as shown by their relative MS intensities. The vertical axis displays the names of the KEGG pathways, and the horizontal axis indicates the number of proteins in each pathway. (C) &(F) Interaction networks of IBV (C) and SARS-CoV-2 (F) nsp3-host cell proteins. The intensity of interactions is represented by the darkness of the color and line thickness, as sourced from STRING. A darker color means a higher number of interacting nodes, and a thicker line between nodes signifies a stronger interaction intensity.

Similar approaches used to analyze SARS-CoV-2 nsp3-interacting proteins showed that proteins with the highest MS intensity were nucleoside diphosphokinase 1 and 2 (NME1/NME2), primarily found in both nucleus and cytoplasm (Figure 3C). Functionally, both NME1 and NME2 can bind to DNA, with NME1 specifically exhibiting a DNA 3’-5’ exonuclease activity. Both proteins play significant roles in DNA repair and regulation of cell transcription as transcription factors [20]. KEGG pathway analysis revealed that 11 out of the top 50 proteins were enriched in the spliceosome pathway (Figure 3D).

Protein-protein interaction (PPI) networks for the top 50 proteins with nsp3 from IBV and SARS-CoV-2, respectively, were then constructed using the STRING database (Figure 3E and F), classifying the interacting proteins with similar functions by high levels of aggregation. Proteins with highest connectivity are GAPDH-HSPA8-NPMI-HNRNPA2B1-HNRNPAl and HNRNPA1-HNRNPA2B1-HNRNPC-SRSF1-GAPDH in the IBV nsp3-host protein and SARS-CoV-2 nsp3-host protein interactomes, respectively (Figure 3E and 3F). As these proteins may be crucial in influencing the host cell metabolism or signal transduction pathways of the entire system, they would be suitable candidates for future functional characterization.

### 3.4 Validation and preliminary characterization of the interactions between seven selected host proteins and nsp3

The accuracy and reliability of the mass spectrum results were then verified by co-immunoprecipitation (CO-IP) assay either in cells overexpressing HiBit-tagged nsp3 and a selection of six potential interacting proteins, or in nsp3-overexpressing cells with an endogenous cellular protein. Six potential interacting proteins, DDX5, DDX39B, DHX9, elF4A3, SRRT, and UBA1, were first selected, and expression plasmids were constructed with a FLAG tag placed at the N-terminus of each protein. These constructs along with nsp3 from IBV and SARS-CoV-2, respectively, were co-transfected into HEK293T cells, samples were collected at 24 h post-transfection, and subjected to CO-IP with the corresponding labeled affinity gel, following by Western blotting with the FLAG or HiBiT antibody. As shown in Figure 4A, we observed detectable bands for both IBV/SARS-CoV-2 nsp3 and one of the co-expressed six proteins in the input group, demonstrating the successful expression of both proteins in HEK293T cells. In the CO-IP experiment with the FLAG affinity gel, the IBV/SARS-CoV-2 nsp3 was detected only when it was co-expressed with one of the selected proteins, confirming that nsp3 from both coronaviruses was indeed interacted with these selected interacting proteins (Figure 4A).

**Figure 4.**
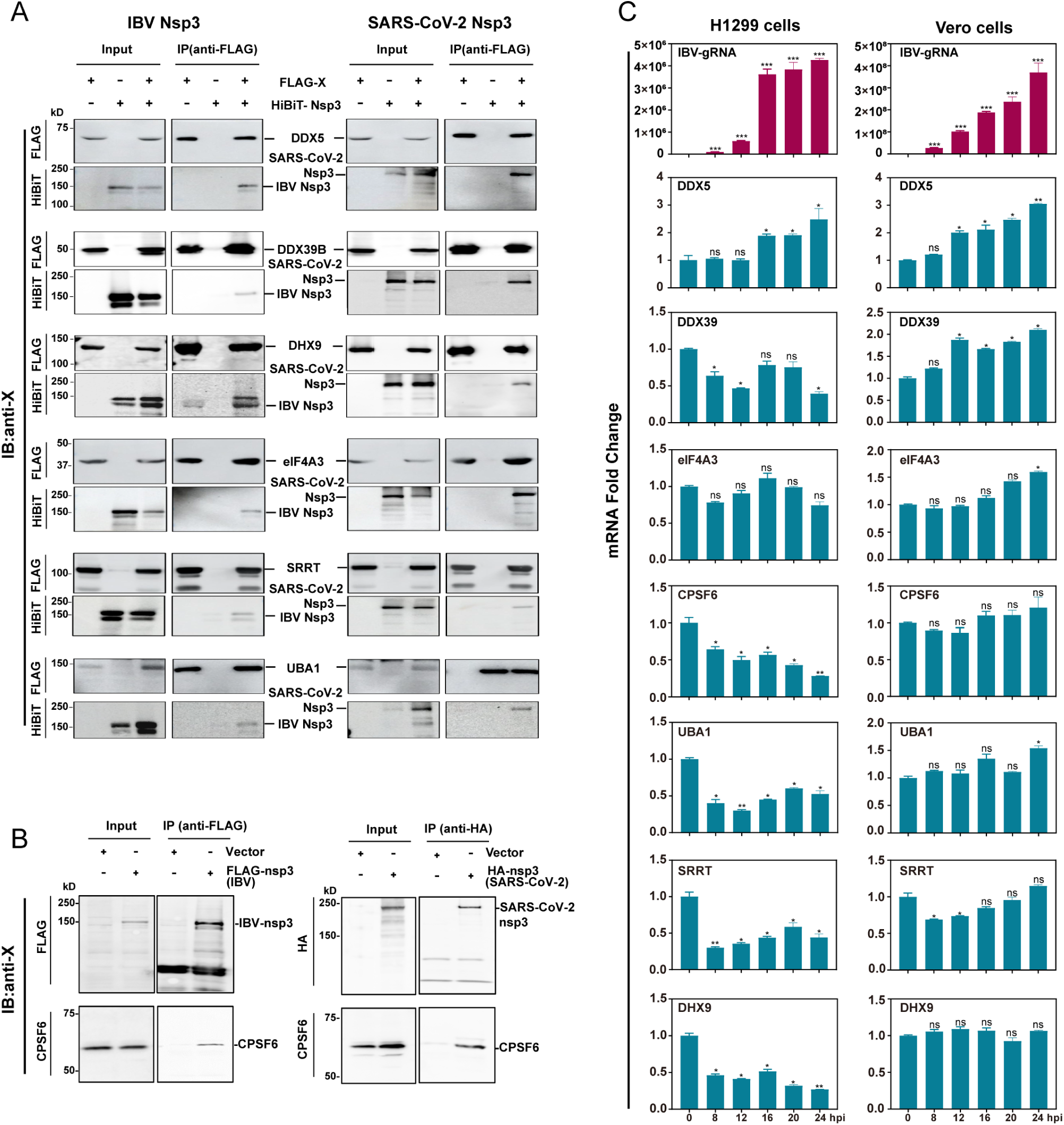
Validation of the interactions between nsp3 and a selection of host proteins. (A) Interaction of IBV and SARS-CoV-2 nsp3 with a selected host protein in cells overexpressing the proteins. HEK293T cells were co-transfected with plasmids encoding HiBiT-tagged IBV or SARS-CoV-2 nsp3, together with a FLAG-tagged host protein (DDX5, DDX39B, DHX9, eIF4A3, SRRT, and UBA1), followed by IP with anti-FLAG beads. The total cell lysates (input) and immunoprecipitated samples were analyzed by immunoblot with anti-FLAG and HiBiT, respectively. The molecular weights of the proteins are indicated on the left and the names of the corresponding proteins are shown on the right. (B) Interaction of endogenous CPSF6 in cells overexpressing either IBV or SARS-CoV-2 nsp3. HEK293T cells were transfected with plasmid encoding FLAG-tagged IBV nsp3 or HA-tagged SARS-CoV-2 nsp3, followed by IP with anti-Flag or HA beads. The total cell lysates (input) and immunoprecipitated samples were analyzed by immunoblot with anti-FLAG and HiBiT, respectively. The molecular weights of the proteins are indicated on the left and the names of the corresponding proteins are shown on the right. (C) Regulation of the transcription of seven selected genes by IBV infection of H1299 and Vero cells. Cells were infected with IBV at a multiplicity of infection (MOI) of approximately 2, and harvested at indicated time points for RNA extraction. Equal amounts of total RNA were reverse-transcribed, and the level of IBV gRNA as well as mRNA levels of DDX5, DDX39B, DHX9, CPSF6, elF4A3, SRRT, and UBA1 were determined by qPCR. Significance levels are presented by the P value (ns, no significance, *, P < 0.05;**, P < 0.01; ***, P < 0.001).

The seventh protein chosen for validation is CPSF6. The interaction between nsp3 and CPSF6 was validated by overexpression of FLAG-tagged IBV nsp3 and HA-tagged SARS-CoV-2 nsp3, respectively, in HEK293T cells, and analyzed by CO-IP. In the input group, the exogenous expression of nsp3 and the endogenous CPSF6 was readily detected with anti-FLAG, anti-HA and anti-CPSF6 antibodies, respectively (Figure 4B). In the CO-IP experiments, CPSF6 was detected only in nsp3-overexpressing cells, but not in control cells transfected with the empty vector (Figure 4B), confirming the interaction of nsp3 from both coronaviruses with the endogenous CPSF6.

As an initial step in characterizing these interactions, we examined the mRNA levels of DDX5, CPSF6, DDX39B, DHX9, elF4A3, SRRT, and UBA1 in cells infected with IBV by time course experiments in H1299 and Vero cells (Figure 4C). Cells were harvested at 0, 8, 12, 16, 20 and 24 h post-infection, respectively, total RNA was extracted and analyzed by RT-qPCR. Among the seven genes, significant upregulation of DDX5 were observed in IBV-infected H1299 and Vero cells, demonstrating that the expression of this gene is induced by IBV infection (Figure 4C). The mRNA levels of DDX39B, DHX9, CPSF6, SRRT, UBA1 and elF4A3 were either unchanged or significantly reduced, but different expression profiles were detected in the two infected cells (Figure 4C). These discrepancies in the expression of these nsp3-interacting proteins in different cell types may reflect different replication kinetics of IBV. It may also suggest that IBV infection may modulate the expression of these genes in different cell types.

### 3.5 Interaction of nsp3 and CPSF6 in IBV-infected cells

As nsp3 interacts with multiple important players in the process of mRNA splicing via spliceosome and translation, we focused on CPSF6 in the subsequent sections. CPSF6, also known as CFIm68, is a 68 kDa component of the mammalian cleavage factor I (CFIm) complex that plays a crucial role in regulating mRNA alternative polyadenylation and determining the length of the 3’ UTR, the two essential mechanisms controlling gene expression [21]. We first set up to confirm if nsp3 was indeed interacted with CPSF6 in virus-infected cells by using rIBV-HA-3541, a recombinant IBV carrying an HA tagged nsp3 as previously described [14]. In H1299 cells infected with rIBV-HA-3541, but not in cells infected with rIBV-p65, the interaction of HA-tagged nsp3 with CPSF6 was clearly detected (Figure 5A).

**Figure 5.**
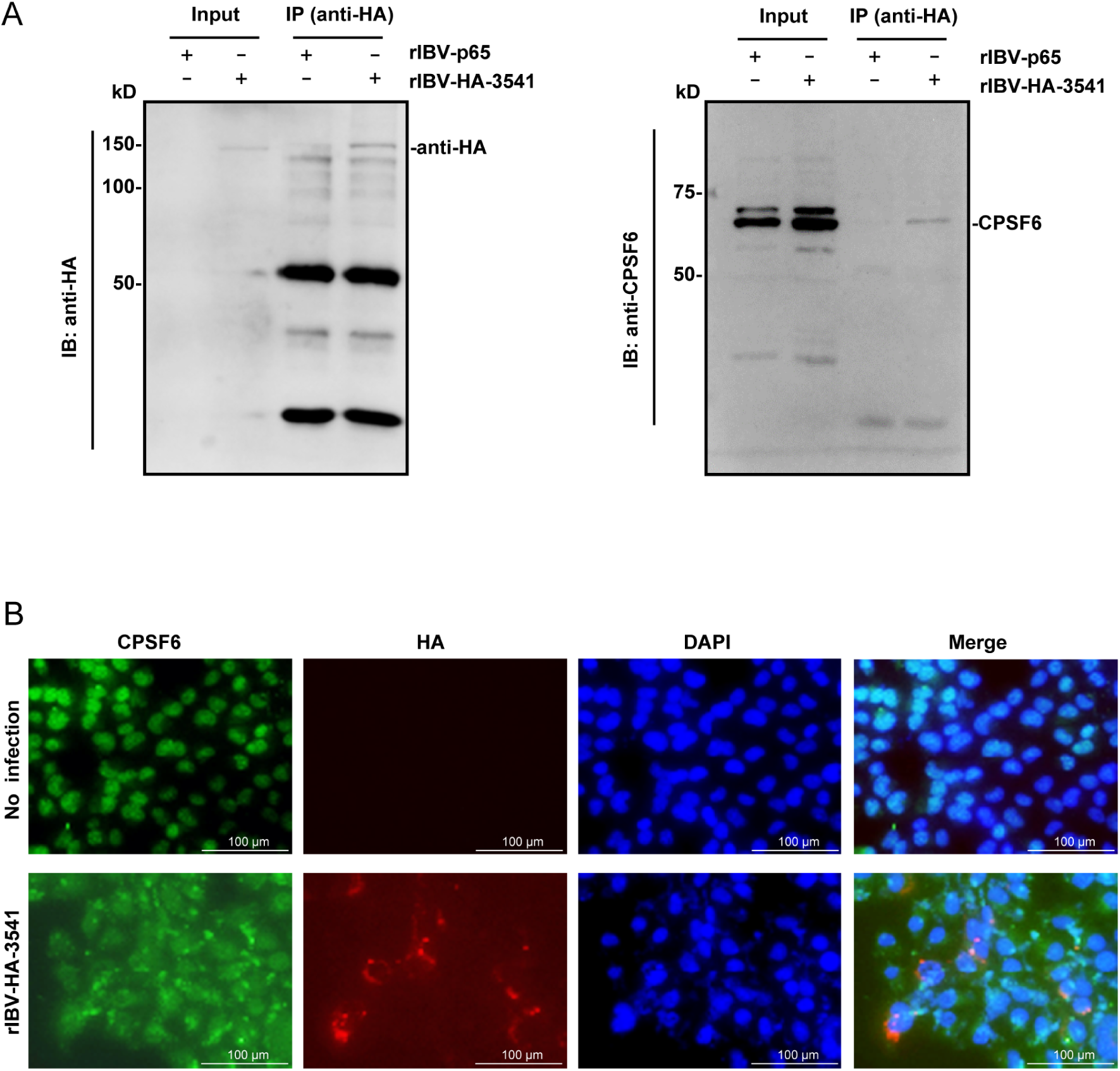
Verification of the interaction between IBV nsp3 and CPSF6 in viral infection. (A) Interaction of IBV nsp3 with CPSF6 in IBV-infected cells. H1299 cells were infected with wild type (rIBV-p65) or a recombinant IBV carrying an HA-tagged nsp3 (rIBV-HA-3541) at an MOI of approximately 2, and harvested 24 h post-infection for protein extraction. The samples were immunoprecipitated with anti-HA beads, followed by immunoblot using anti-HA and anti-CPSF6 antibodies. The total lysates as included as the input. The molecular weights of the proteins are indicated on the left and the names of the corresponding proteins are shown on the right. (B) Cytoplasmic translocation and co-localization of CPSF6 with nsp3 in IBV-infected H1299 cells. Cells were infected with rIBV-HA-3541 at an MOI of approximately 2, and fixed at 24 hpi. Immunofluorescent staining was performed using anti-CPSF6 and HA antibodies, and nuclear stained with DAPI. The fluorescence signals were examined and images taken by fluorescence microscopy (x200). The merged images show CFSP6 and IBV nsp3 colocalization.

As this interaction may change the subcellular localization of CPSF6 in IBV-infected cells, H1299 cells were then infected with rIBV-HA-3541, fixed at various time points, and immunostained with anti-CPSF6 and anti-HA antibodies, followed by the application of fluorescent secondary antibodies for visualization. CPSF6 was primarily concentrated in the nucleus of uninfected cells. In contrast, a certain proportion of CPSF6 was relocated to the cytoplasm and colocalized with IBV nsp3 in the infected cells (Figure 5B). These results demonstrated that the interaction between IBV nsp3 and CPSF6 alters the subcellular localization of CPSF6, suggesting that the cytoplasm-translocated CPSF6 may facilitate viral genome synthesis through its interaction with nsp3.

### 3.6 Promotion of IBV replication by nsp3-CPSF6 interaction

To study the functional role of CPSF6 in viral replication, we first designed siRNA targeting the CPSF6 sequence to knock down its expression in H1299 cells before infection with rIBV-Hibit, a recombinant IBV with a Hibit tag inserted in the IBV structural protein S using a reverse genetics system [17]. It allows to accurately determine the virus replication efficiency and viral titers through a rapid chemiluminescence method, as previously described [17]. The supernatants and cells were harvested for analysis of proteins and RNA at 16, 24, and 32 h post-infection, respectively, and Western blot analysis was carried out, demonstrating efficient knockdown of CPSF6 in cells transfected with siCPSF6, compared with siEGFP-transfected control cells (Figure 6A). In the CPSF6-knockdown cells infected with IBV, generally lower levels of IBV structural proteins N, S, and M than those in the control cells transfected with siEGFP were detected, with particularly lower levels of IBV N and S proteins detected at 16 hpi (Figure 6A). RT-qPCR analysis of gRNA, -gRNA and sgRNA showed marked reduction of these viral RNAs in the siCPSF6-knockdown cells infected with IBV, compared with those in the siEGFP-control cells infected with IBV (Figure 6B). Consistently, luminescence measurement showed much lower levels of fluorescence values in the CPSF6-knockdown cells infected with IBV than those in the control cells infected with the same virus (Figure 6C).

**Figure 6.**
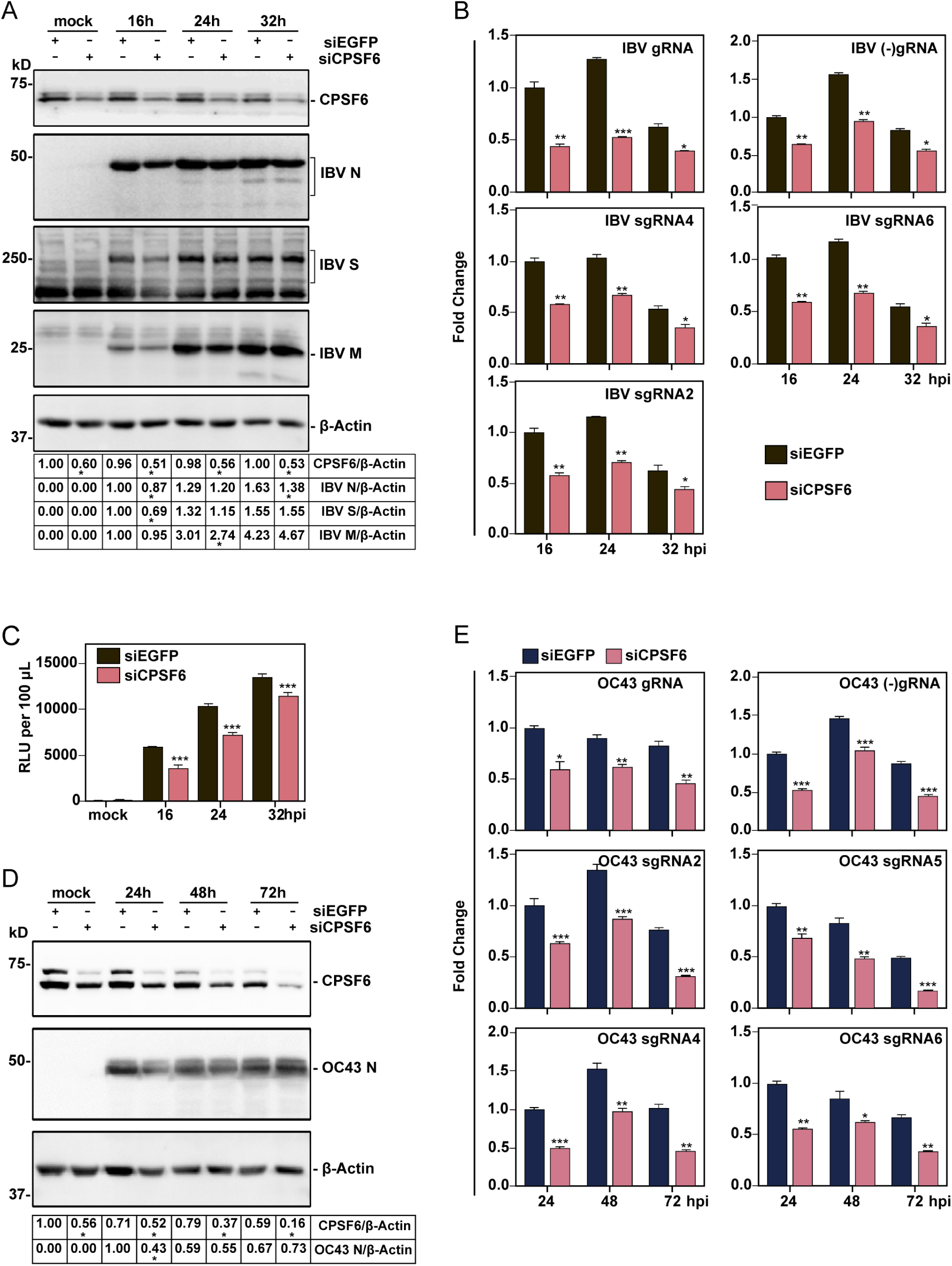
Knockdown of CPSF6 inhibits the replication of IBV and HCoV-OC43. (A) Western blot analysis of the effect of CPSF6-knockdown on the replication of IBV. H1299 cells were transfected with siEGFP and siCPSF6 before being infected with IBV at an MOI of 2. Cell lysates were harvested at the indicated time points and subjected to Western blot analysis using the indicated antibodies. Beta-actin was included as the loading control. Protein ladders in kilodaltons are indicated on the left. Significance levels are presented by the P value (*, P<0.05) (B) RT-qPCR analysis of the effect of CPSF6-knockdown on the replication of IBV. Total RNAs were extracted from cells infected and harvested as described in panel (A). Equal amounts of total RNA were reverse-transcribed, and the level of IBV gRNA, -gRNA and sgRNAs were determined by qPCR. Significance levels are presented by the P value (*, P < 0.05;**, P < 0.01; ***, P < 0.001). (C) HiBiT luminescence assessment of the effect of CPSF6-knockdown on the replication of IBV. Cells were infected as described above in panel (A), and the culture supernatants were collected at the specified time point for HiBiT luminescence. Significance levels are presented by the P value (***, P < 0.001) (D) Western blot analysis of the effect of CPSF6-knockdown on the replication of HCoV-OC43. H1299 cells were transfected with siEGFP and siCPSF6 before being infected with HCoV-OC43 at an MOI of 2. Cell lysates were harvested at the indicated time points and subjected to Western blot analysis using the indicated antibodies. Beta-actin was included as the loading control. Sizes of protein ladders in kilodaltons are indicated on the left. Significance levels are presented by the P value (*, P<0.05) (E) RT-qPCR analysis of the effect of CPSF6-knockdown on the replication of HCoV-OC43. Cells were treated as described above in panel (D), and harvested at indicated time points for RNA extraction. Equal amounts of total RNA were reverse-transcribed. The level of OC43 gRNA and sgRNAs were determined by qPCR. Significance levels are presented by the P value (*, P < 0.05;**, P < 0.01; ***, P < 0.001)

The effect of CPSF6 knockdown on the replication of another coronavirus was then carried out similarly in CPSF6-knockdown H1299 cells infected with HCoV-OC43. Cells were harvested at 24, 48, and 72 hpi, and analyzed by Western blot and RT-qPCR. Efficient knockdown of CPSF6, once again, was observed in cells transfected with siCPSF6, compared to that in the control cells transfected with siEGFP (Figure 6D). The expression of HCoV-OC43 structural protein N was significantly reduced in CPSF6-knockdown cells at 24 hpi, compared with that in the control cells (Figure 6D). Decreased levels of gRNA, -gRNA and sgRNA, similar to those observed in IBV infected cells, were also detected in CPSF6-knockdown cells (Figure 6E). These results suggest that CPSF6 plays an important role in promoting viral replication in infected cells, potentially linked to the replication of viral gRNA and sgRNA.

### 3.7 Modulation of sgRNA transcription by nsp3-CPSF6 interaction

As CPSF6 would impose a more prefunded impact on the selection and transcription of sgRNAs located more closely to the 3’ end of the genome, the effect of CPSF6-knockdown on the transcription of these sgRNA were studied by RT-PCR amplification and sequencing using a strategy outlined in Figure 7A. RT-PCR amplification of the two recently identified IBV sgRNAs [22,23]. referred to as 3’-noncoding sgRNA1 and 2 in this study, and sgRNA6 coding for IBV N protein using a primer pair spanning the leader region and the end of the 3’-UTR would generate three fragments with sizes of 212, 497 and 1893 bp, respectively (Figure 1A). Accordingly, we collected samples from IBV-infected CPSF6-knockdown cells and performed Western blot and RT-PCR analyses. Once again, the efficient knockdown of CPSF6 was observed in cells transfected with siCPSF6, compared with that in siEGFP-transfected control cells (Figure 7B). Gel electrophoresis analysis of RT-PCR-amplified fragments showed a significant reduction in the sgRNA6 band and nearly undetectable bands for 3’-noncoding sgRNA1 and 2 in siCPSF6-knockdown cells infected with IBV (Figure 7C). The amplified fragments were further analyzed by cloning into a vector and sequencing, confirming that these two amplified fragments were indeed representing the two small noncoding sgRNA (Figure 7D). These results demonstrate that translocation of CPSF6 to the cytoplasm and clustering to the viral RNA replication sites by its interaction with nsp3 play a positive regulatory role in the transcription of IBV sgRNAs close to the 3’UTR of the genome.

**Figure 7.**
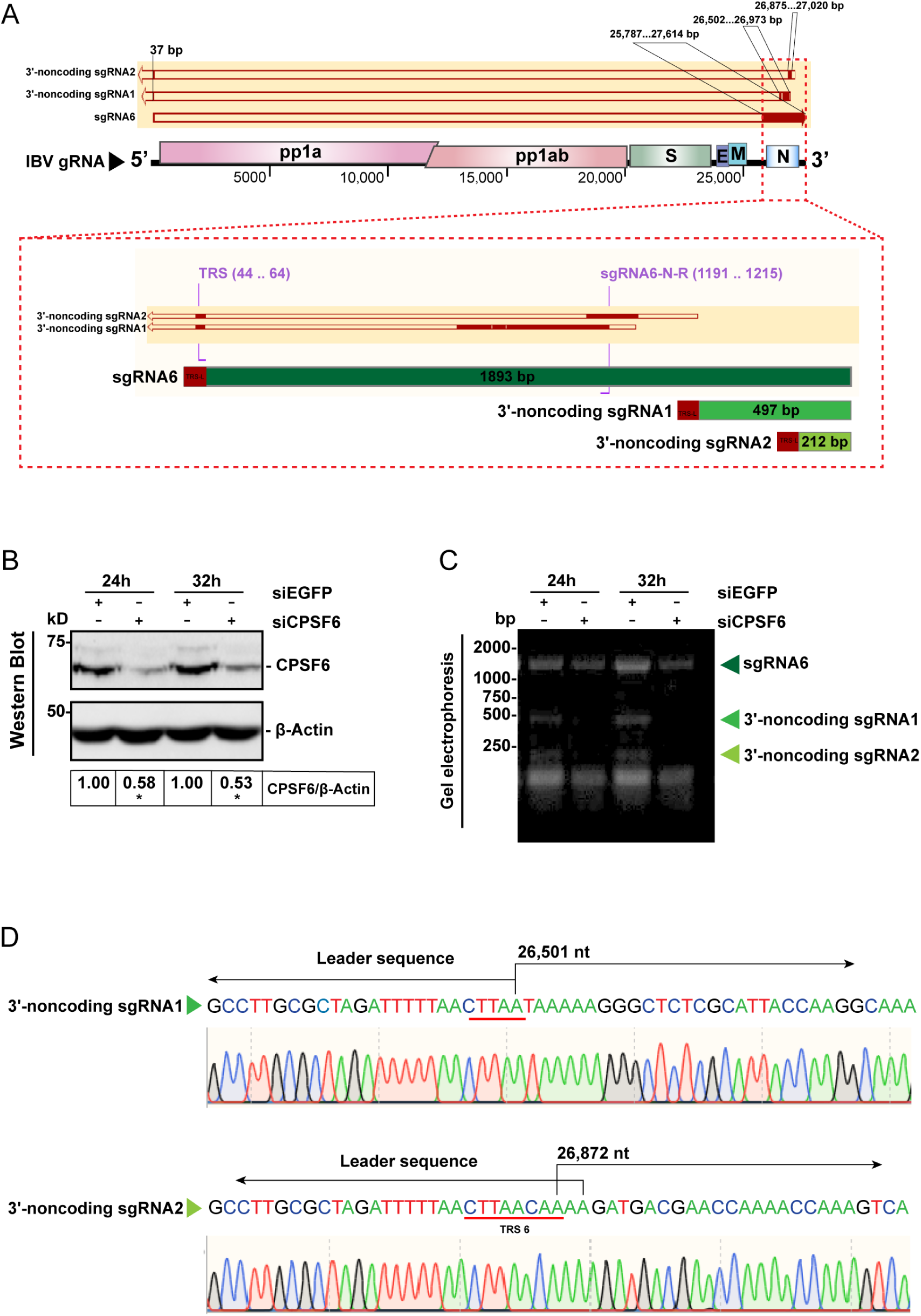
Effects of nsp3-CPSF6 interaction on viral sgRNA transcription. (A) Diagram showing the location of three sgRNAs at the 3’-region of the genome of IBV. Shown are the lengths in nucleotides of sgRNA6, 3’-noncoding sgRNA1 and 2. (B) Analysis of the CPSF6 knockdown efficiency and its effect on IBV replication by Western blot. Cells were transfected with siEGFP and siCPSF6 before being infected with IBV at an MOI of 2, harvested at the indicated time points, lysates prepared and subjected to Western blot analysis using the indicated antibodies. Sizes of proteins are indicated on the left. Significance levels are presented by the P value (*, P<0.05) (C) The effect of CPSF6-knockdown on the replication of sgRNAs located at the 3’ end region of the IBV genome. Cells were treated as (B), total RNAs were extracted and subjected to RT-PCR analysis, using the primer pair TRS-L-F and sgRNA6-N-R. Sizes of PCR products are indicated on the left. (D) Confirmation of PCR amplified fragments by nucleotide sequencing. The sequences at junctions between IBV leader at the 5’-UTR and the bodies of the two noncoding sgRNAs at the 3’-UTR of the viral genome confirm the identifies of the PCR-amplified fragments described in (C) and the effect of CPSF6-knockdown on the transcription of these two sgRNAs.

## 4. Discussion

As the largest coronavirus protein, nsp3 encodes multiple functional domains and plays crucial roles in viral replication and immune evasion [4]. This huge size and multifunctional nature would predestine the formation of complex interaction networks with host cell factors to modulate its viral and cellular functions, and vice versa. However, the large size, complex structural features and the presence of multiple transmembrane domains as well as potential toxic regions to bacteria make cloning and efficient expression of the full-length nsp3 proteins from different coronaviruses challenging, resulting in the relative scarcity of reports on the interactomes of nsp3 and host cell factors. In this study, we report the application of a two-plasmid system employing K1E phage RNA polymerase along with a potent promoter to efficiently and accurately express the full-length nsp3 from two distantly related coronaviruses, gammacoronavirus IBV and betacoronavirus SARS-CoV-2 [14], to map and comparatively study the nsp3-host protein interactomes. Our findings revealed that 1,150 host proteins interacted with IBV nsp3, while 920 host proteins interacted with SARS-CoV-2 nsp3, with 658 common interacting host proteins sharing by the two nsp3 proteins. Further validation and preliminary characterization of selected candidates demonstrated the reproducibility and reliability of the proteomics data and revealed an essential function of nsp3-CPSF6 interaction in regulation of coronavirus sgRNA transcription (Figure 8).

**Figure 8.**
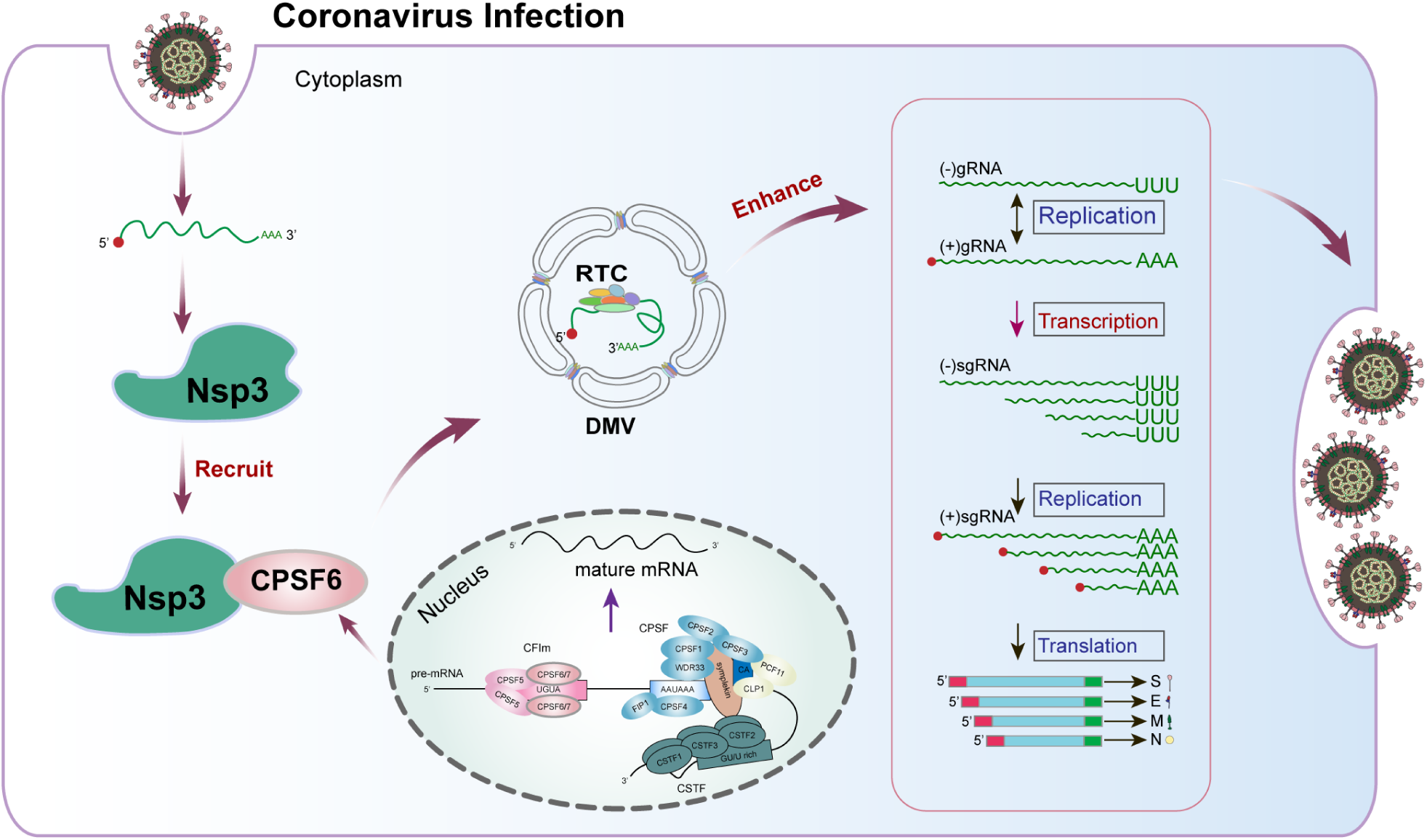
Diagram illustrating the current working model. Illustrated are our current working model. Coronavirus nsp3 may recruit a certain proportion of CPSF6 to the cytoplasm and cluster to the viral replication sites, facilitating the replication and transcription of viral gRNA and sgRNAs, especially sgRNAs located at the 3’-region of the genome. RTC, replication/transcription complex; DMV, double-membrane vesicles.

The demonstration that host proteins interacting with nsp3 from these two distantly related coronaviruses shared high similarities in terms of subcellular localization, functions, and the pathways involved, particularly those associated with RNA metabolism and protein synthesis. Similar observations were also documented in a previous study using nsp3 from SARS-CoV and SARS-CoV-2, presenting multi-omics data, including interactome, phosphoproteome, ubiquitylome, transcriptome and proteome. As the full-length nsp3 from these two closely related coronaviruses share much higher amino acid identity (∼76%) than that from the two distantly related coronaviruses (IBV and SARS-CoV-2, ∼20%) presented in this study, the observation that similar host cell factors associated with RNA metabolism and protein synthesis would reveal a general involvement of these cellular factors and pathways in the coronavirus replication cycle. On the other hand, the two studies have also identified many distinct host proteins that potentially interact with individual nsp3 from different coronaviruses. In additional to the general discrepancies and poor reproducibility of the omics data from different studies, it may reflect the involvement of specific host cell factors in the replication and pathogenicity of different coronaviruses. The application of a unique cytoplasmic and highly efficient expression system in this study has provided a nsp3 preparation with a majority of the expressed nsp3 protein as full-length [14], and would ensure a high quality and reliability of the interaction networks constructed. This was, in fact, supported by the subsequent validation of the seven randomly selected candidate interactors presented in this study. As the top 50 nsp3-interacting host proteins were enriched in the spliceosome pathway, particularly for the SARS-CoV-2 nsp3, the verified interactions of these seven candidates associated with the spliceosomal and RNA transport pathway reinforced the conclusion that coronavirus nsp3 may recruit these host factors to regulate viral RNA synthesis, ultimately promoting viral replication and pathogenesis.

DDX5, eIF4A3, DDX39B, and DHX9 belong to the DExD/H box protein family and play crucial roles in RNA metabolism [24], and are widely reported to interact with various viral proteins. DDX5 was shown to interact with a variety of proteins from RNA viruses to promote viral replication, including SARS-CoV helicase nsp13 [25] and SARS-CoV-2 N protein [26]. Other examples include its interaction with the Rev and Tat of human immunodeficient virus-1 (HIV-1) to act as a co-factor for both regulation of HIV-1 RNA transcription and enhancement of the nuclear export of HIV-1 transcripts [27,28], and its interaction with hepatitis C virus (HCV) NS5B allows recruitment of the protein into the viral replication complex and enhancement of HCV replication [29]. DDX5 also acts as a positive regulator for the replication of Japanese encephalitis virus (JEV) and the porcine reproductive and respiratory syndrome virus (PRRSV) through its interactions with proteins from these viruses [30,31]. eIF4A3 was reported to facilitate the formation of spliced viral mRNA and its nuclear export in the life cycles of human cytomegalovirus (HCMV) [32] and herpes simplex virus (HSV) [33], as well as to interact with influenza A virus (IAV) PB2 protein [34]. DDX39B (UAP56) was reported to interact with multiple proteins from SARS-CoV-2, including NSP5, NSP6, NSP13, NSP14, and S proteins [35]. Recently, DDX39B and its paralog DDX39A (URH49) were certified to exert a dual effect on regulating the polymerase activity and replication of IAV by interacting with viral NP and NS1 proteins [36–39]. DHX9 contributes to various aspects of viral function, enhancing the infectivity of multiple viruses, including HIV-1, HCV, IAV, and classical swine fever virus [40]. Interestingly, DHX9 also possesses the antiviral activity in Epstein-Barr virus (EBV) infection by interacting with EBV essential protein SM and restricting EBV lytic replication [41].

So far, the association of SRRT and UBV1 with viral proteins is less reported. SRRT, also referred to as Ars2, plays a vital role in miRNA biogenesis and is crucial for cell proliferation. It may link the nuclear cap-binding complex to RNA interference and cell proliferation, as depletion of Ars2 significantly reduces pri-miRNA processing and the levels of several miRNAs implicated in transformation, including miR-21, let-7, and miR-155 [42]. UBA1 is a key regulator of protein homeostasis and functions as an E1 ubiquitin-activating enzyme, playing a crucial role in the regulation of cellular protein homeostasis [43]. A new study has revealed the UBA1-STUB1 axis mediates cancer immune escape and resistance to checkpoint blockade [44].

CPSF6 is a crucial component of the CFIm complex, modulating mRNA alternative polyadenylation (APA) and maturation of pre-mRNA to functional mRNA [21]. This protein is also involved in viral mRNA processing. Interaction of CPSF6 with the NP1 proteins from minute virus of canines (MVC) and human bocavirus 1 (HBoV1) was shown to regulate viral alternative RNA splicing, modulate viral mRNA processing and viral DNA replication [45]. Knockout of CPSF6 was also reported to reduce HIV-1 transcription by inhibiting RNA polymerase II (Pol II) and cyclin-dependent kinase 9 (CDK9) phosphorylation [46]. The demonstration in this study that knockdown of CPSF6 suppressed IBV replication, particularly the transcription of sgRNAs located at the 3’ end of the genome may reveal a novel mechanism exploited by coronavirus to facilitate the replication of its gRNA and sgRNA.

Transcription and replication of coronavirus gRNA and sgRNA are crucial for successful completion of the viral replication cycle. These processes are extensively regulated by both viral and host factors. Earlier studies have focused on the roles of viral nsps and RNA elements, highlighting the key regulatory functions of leader TRS (TRS-L) and body TRS (TRS-B) in sgRNA transcription. The essential regulatory role of the complementarity between TRS-L and TRS-B in sgRNA transcription was underscored in the formation of the two newly identified sgRNAs and in our previous study that alteration of certain bases in TRS-B using an IBV infectious cloning system significantly reduces or completely inhibits the transcription of the corresponding sgRNA [22,23]. In terms of host proteins involved in the regulation of these processes, various genome-wide screening approaches have been used to identify and characterize host proteins, including the identification and characterization of members of the heterogeneous nuclear ribonucleoprotein (hnRNP) family and MADP1 interacting with gRNA [2,47], and DDX1 interacting with nsp14 [48]. More recently, several host proteins, including T-cell-restricted intracellular antigen-1 (TIA1), Insulin Like Growth Factor 2 MRNA Binding Protein 1 (IGF2BP1), Polypyrimidine Tract Binding Protein 1 (PTBP1), have been identified to interact with the SARS-CoV-2 RNA genome, and a comprehensive landscape of the SARS-CoV-2 ncrRNA–host protein interactome has been depicted [49–51]. However, the mechanisms underlying the regulatory roles of those factors on viral sgRNA transcription remain largely unknown. In this study, we report the crucial role of IBV nsp3-CPSF6 interaction in the transcription of small noncoding sgRNAs located at the 3’UTR of the IBV genome, pointing out the important involvement of host RNA splicing and polyadenylation machineries in coronavirus RNA replication and transcription.

In conclusion, we have constructed the interactomes of nsp3 from IBV and SARS-CoV-2 with host cell factors, demonstrating that multiple host cell proteins and important cellular pathways are shared by the two coronavirus nsp3 proteins. Furthermore, an essential role of nsp3-CPSF6 interaction in regulation of coronavirus gRNA and sgRNA transcription has been revealed. These findings have provided new information on our current understanding of host cell factors in coronavirus replication and a framework for further functional studies.

## CRediT authorship contribution statement

Xinxin Sun: Conceptualization, Data curation, Methodology, Investigation, Visualization, Formal analysis. Lixia Yuan: Data curation, Methodology, Investigation, Visualization, Formal analysis, Writing & review and editing, Funding acquisition. Zihong Hu: Methodology, Investigation, Writing . Yuzhu Lai: Data curation, Investigation, Validation, Visualization. Bei Yang: Investigation, Validation, Visualization. Jiawen He: Data curation, Investigation, Validation, Visualization. Ruiai Chen: Investigation, Validation, Visualization. Ding Xiang Liu: Conceptualization, Funding acquisition, Supervision, Writing-review and editing.

## Acknowledgments

This work was partially supported by the National Natural Science Foundation of China (grant number 32170152), Guangdong Basic and Applied Basic Research Foundation of China (grant number 2024A1515012930), Guangdong Provincial Laboratories Research and Innovation Programs (grant number XJGDL2022001), Zhaoqing Xijiang Innovative Team Foundation of China (grant numbers P20211154-0201 and P20211154-0202,), and Open Research Project of the Key Laboratory of Viral Pathogenesis & Infection Prevention and Control of the Ministry of Education (2024VPPC-R05).

